# Juvenile Huntington’s Disease and Other PolyQ diseases, Update on Neurodevelopmental Character and Comparative Bioinformatic Review of Transcriptomic Data

**DOI:** 10.1101/2021.02.19.431958

**Authors:** Karolina Świtońska-Kurkowska, Bart Krist, Joanna Maria Delimata, Maciej Figiel

## Abstract

Polyglutamine (PolyQ) diseases are neurodegenerative disorders caused by the CAG repeat expansion mutation in affected genes resulting in toxic proteins containing a long chain of glutamines. There are nine PolyQ diseases: Huntington’s disease (HD), spinocerebellar ataxias (types 1, 2, 3, 6, 7, and 17), dentatorubral-pallidoluysian atrophy (DRPLA), and spinal bulbar muscular atrophy (SBMA). In general, longer CAG expansions and longer glutamine tracts lead to earlier disease presentations in PolyQ patients. Rarely, cases of extremely long expansions are identified for PolyQ diseases, and they consistently lead to juvenile or sometimes very severe infantile-onset polyQ syndromes. In apparent contrast to the very long CAG tracts, shorter CAGs and PolyQs in proteins seems to be the evolutionary factor enhancing human cognition. Therefore, polyQ tracts in proteins can be modifiers of brain development and disease drivers, which contribute neurodevelopmental phenotypes in juvenile- and adult-onset PolyQ diseases. Therefore we performed a bioinformatics review of published RNAseq polyQ expression data resulting from the presence of polyQ genes in search of neurodevelopmental expression patterns and comparison between diseases. The expression data were collected from cell types reflecting stages of development such as iPSC, neuronal stem cell, neurons, but also the adult patients and models for PolyQ disease. Our comparative bioinformatic review highlighted several (neuro)developmental pathways and genes identified within PolyQ diseases and mouse models responsible for neural growth, synaptogenesis, and synaptic plasticity.

## 1. PolyQ diseases and Juvenile Cases

Polyglutamine (PolyQ) diseases are neurodegenerative disorders caused by expansion mutations giving rise to abnormally long CAG tri-nucleotide repeat tracts in affected, otherwise unrelated genes. PolyQ disorders are dominantly inherited and autosomal, except for SBMA, which is X-linked. The expanded polyQ repeats disturb the function of the proteins encoded by the genes with CAG expansion, leading to loss or gain of function (Lim et al., 2008). To date, nine PolyQ diseases were identified; namely Huntington’s disease (HD), spinocerebellar ataxia (SCA) types 1, 2, 3, 6, 7, and 17, dentatorubral-pallidoluysian atrophy (DRPLA), and spinal bulbar muscular atrophy (SBMA) (Zoghbi and Orr, 2009).

Several PolyQ diseases may occur in younger patients, and in such cases, symptom presentation in juvenile disease usually differs from the adult form. Although the juvenile and infantile forms make up a minority of instances, the early onset, and polyQ protein domains, usually much longer than in the adult forms, hint at the developmental nature of these cases. Since the etiology of the diseases is genetic and more defined, they may help to better understand the brain development in health and disease. The first aim of this work is to obtain a broader literature overview of the juvenile and infantile PolyQ disease cases with very long CAG repeats in the context of early brain development. Since brain development is primarily related to the forming of new cell populations, differentiation, and wiring of the brain, we also looked at what is known about these processes in the context of juvenile polyQ cases. In the second part of the work, we performed a bioinformatics analysis of RNAseq data from polyQ patients and models in search of neurodevelopmental expression patterns and comparison between diseases. The expression data were collected from cell types reflecting stages of development such as iPSC, neuronal stem cell, neurons, but also the adult patients and models for PolyQ disease. In addition, thanks to a broader selection of transcriptomic data in mice containing longer CAG tracts, we are able to compare gene expression profiles between different PolyQ diseases. Still, the bias towards HD in this work results from the available data sources. However, another aim of our work is the focus on juvenile cases of polyQ disorders other than HD, possible neurodevelopmental signs in the diseases, and what we could still learn from the juvenile forms about diseased brain development.

### 1. Juvenile and Infantile Huntington’s Disease

In HD, the CAG expansion mutation is located in the Huntingtin (*HTT*) gene (The Huntington’s Disease Collaborative Research Group, 1993), which is crucial for neural development (reviewed in (Saudou and Humbert, 2016)). The juvenile form of HD (Juvenile onset Huntington’s disease; JOHD) is defined as disease onset before the age of 20 with the number of CAG repeats between 60 (Quarrell et al., 2013) and 89 (Nance and Myers, 2001; Ribaï et al., 2007), and infantile HD with very rapid onset with number of CAG repeats above 90 and more (Fusilli et al., 2018; Stout, 2018). JOHD is also marked by a more rapid disease progression, leading to an earlier death (Fusilli et al., 2018). In JOHD, the symptoms are typically seizures, rigidity, and severe cognitive dysfunction (Nance and Myers, 2001; Vargas et al., 2003; Squitieri et al., 2006; Ribaï et al., 2007). In cases where the onset is very early (before ten years of age, sometimes also referred to as “infantile-” or “ultra-juvenile HD”), epilepsy is also frequent (Barbeau, 1970). One of the youngest onset of JOHD and also one of the most severe presentation which has been described to date was a girl who had healthy development until 18 months of age and later at the age of 3,5 years, showed marked cerebellar atrophy. Around the age of four, choreiform movements on the right side developed. The patient was diagnosed to have 265 triplet repeats on the mutant *HTT* allele and 14 on the other (Milunsky et al. 2003). Other reports have described frequent speech difficulties as early symptoms before motor problems arise (Yoon et al., 2006; Sakazume et al., 2009). Behavioral problems, such as aggression, irritability, and hyperactivity, which are often reported signs of disturbed brain development were also reported for juvenile HD (Yoon et al., 2006).

### 1. Early Onset in PolyQ Spinocerebellar Ataxias

Most severe cases of juvenile or infantile-onset were reported for SCA2 (*ATXN2* gene; ataxin-2 protein), SCA7 (ATXN7; Ataxin-7 protein), SCA17 (TBP gene; TATA Binding protein), and DRPLA (ATN1 gene; atrophin-1 protein). Juvenile-onsets were also reported for SCA3/MJD with more severe presentation compared to adult forms. One of the reasons for the occurrence of very severe developmental signs may be the function of ATXN2, ATXN7, TBP, and ATN1, which can be summarized as a very pleiotropic and broad influence on transcriptional regulation. The function of the genes, including their impact on transcription, has been well-reviewed previously (Shen and Peterson, 2009; Yang et al., 2016; Lee et al., 2018b; Niewiadomska-Cimicka and Trottier, 2019). In SCA1 (*ATXN1* gene; Ataxin-1 protein), SCA3/MJD (*ATXN3* gene; ataxin-3), and SCA6 (*CACNA1A* gene; α1A subunit of the voltage-gated P/Q type channel), the cases with the earliest reported onset were mostly showing signs shortly before adolescence.

The expansion mutation in ATXN2 in infantile cases can be very severe, reaching the range of 124 and 750 CAGs, and the range between 62 and 92 defines onset in early childhood. Typically, SCA2 presents with progressive involuntary movements of the limbs, sensorimotor neuropathy, and slowed eye movements. The abnormal eye movements and myoclonic jerks are generally the first symptoms seen in infantile and early childhood cases, with the onset of disease as early as two months of age (Moretti et al., 2004; Vinther-Jensen et al., 2013; Singh et al., 2014; Sánchez-Corona et al., 2020). Besides these, pigmentary retinopathy, seizures, dysphagia, and early death are unfortunately also standard features of juvenile SCA2 (Babovic-Vuksanovic et al., 1998; Mao et al., 2002).

Abnormally long polyQ tract in the ataxin 7 (*ATXN7*) gene primarily manifests as cerebellar ataxia in SCA7; however, the unique symptom is retinal degeneration, which often is the first presenting symptom (Niewiadomska-Cimicka and Trottier, 2019). In the case of ATXN7, healthy alleles of this gene bear up to 35 CAG repeats, whereas SCA7 affected individuals have more than 39 repeats (David et al., 1997; Stevanin et al., 1998). The childhood-onset of SCA7 is the consequence of more than 100 CAG repeats in the *ATXN7* gene (La Spada, 2020). Besides the classic symptoms of progressive cerebellar ataxia and retinal degeneration, the juvenile cases of SCA7 presented with absent or depressed deep tendon reflexes, which is not the case in the adult-onset type of the disease (Enevoldson et al., 1994). Other studies reported symptoms such as seizures, dysphagia, myoclonus, head lag, the absence of cough reflex, and severe hypotonia, but also symptoms more uncommon for PolyQ diseases such as cardiac involvement, hepatomegaly, multiple hemangiomas, atrial septum defect, patent ductus arteriosus, and congestive heart failure accompany ataxia (Benton et al., 1998a; Johansson et al., 1998; van de Warrenburg et al., 2001; Ansorge et al., 2004). Summarizing, infantile or early childhood SCA7 is a severe developmental syndrome with patient death reported as early as six weeks of age from unspecified cardiac and other anomalies (Neetens et al., 1990).

In DRPLA, the affected gene is *ATN1* (Koide et al., 1994), a transcriptional regulator involved in the brain and other organ development (Palmer et al., 2019). In the case of the *ATN1* gene, CAG repeat sizes can vary between 6 and 35 in healthy individuals, while the expansion of more than 48 repeats results in full penetrance and gives rise to the disease (Nagafuchi et al., 1994). Patients with juvenile-onset DRPLA often have progressive myoclonic epilepsy as one of the first symptoms (Tomoda et al., 1991) and the onset in first years of life with CAG repeats between 70 and 80 (Veneziano and Frontali, 1993; Hasegawa et al., 2010). Disease onset could occur as early as six months of age (with an extreme number of CAG repeats of 90 and 93), when hyperkinetic and involuntary movements, the difficulty of controlling head movements, and seizures developed (Shimojo et al., 2001).

SCA17 is caused by an abnormal number (more than 45-47) of CAG or CAA repeats in the TATA box-binding protein (TBP) (Gao et al., 2008; Toyoshima and Takahashi, 2018). In SCA17, a small gain in CAG number in the TBP gene results in a very severe level of genetic anticipation (Maltecca et al., 2003; Rasmussen et al., 2007). For instance, CAG repeats in the range of 55-58 may cause the disease onset at age 20, 61 CAG was associated with onset at age 11, while 66 CAGs resulted in onset at the age of 3 years. (Koide et al., 1999; Maltecca et al., 2003; Rasmussen et al., 2007). Common features of the disease are ataxic gait, dysarthria, loss of muscle control, seizures, spasticity, tremor, and intellectual dissability. Given the strong anticipation resulting from only low intergenerational expansion, SCA17 and TBP may strongly influence the brain development and transcriptional control of developmental genes.

SCA3 early childhood-onset, described in 2016, involved the range of CAG repeat between 80 and 91 (Donis et al., 2016). The progression of the disease was faster compared to adolescent cases and the signs observed were ataxia, pyramidal findings, and dystonia. In previous SCA3/MJD cohorts, the maximal number of CAGs was 86 (Todd and Paulson, 2010; Tezenas du Montcel et al., 2014).

SCA6 is caused by a polyQ mutation in the calcium channel gene *CACNA1A* (Zhuchenko et al., 1997). SCA6 develops due to a relatively low number of CAG repeats, with 5 to 20 repeats being considered healthy and 21 repeats and above giving rise to the disease (Ishikawa et al., 1997).

The length of CAG repeats in infantile or childhood PolyQ diseases highly influences the onset and severity of the disease. Moreover, genetic anticipation, earlier (and more severe) disease onset in successive generations, is playing a crucial role in the majority of these disorders. (Jones et al., 2017).

SBMA, also referred to as Kennedy disease, is a form of spinal muscular atrophy that is recessive and X-linked, and therefore only occurs in males. The cause of SBMA is a CAG repeat expansion in exon 1 of the androgen receptor gene. Juvenile onset commonly presents with limb atrophy and gynecomastia between 8 to 15 years of age (Echaniz-Laguna et al. 2005). Unlike in other PolyQ diseases discussed here, the number of CAG repeats only poorly predicts the age of onset (muscle weakness) (Sperfeld et al. 2002; Echaniz-Laguna et al. 2005). Because of a more muscle-connected character of a disease, we do not mention it in further paragraphs. However, we included SBMA transcriptomic data in the comparative bioinformatic study.

## 2. Early Brain Development in Health and PolyQ disease

Normal brain development consists of cellular processes such as cell division, cell migration, cell differentiation, maturation, synaptogenesis, and apoptosis, which are precisely orchestrated by a molecular network of signaling pathways. Such orchestration is crucial for the correct generation of cellular layers, specialized neural regions, and the generation of complex neuronal wiring between brain structures. In brief, during the formation of the neural tube (neurulation) in the embryo, the neuroepithelial cells (NECs) perform symmetric cell divisions producing progenitors of different brain regions (Paridaen and Huttner, 2014). Pax6 and Emx2 signaling molecules expressed in opposing gradients from the anterior to posterior regions of the proliferative zone function as a primitive blueprint for the dividing NECs to give rise to the early structures of the forebrain, midbrain, and hindbrain (Stiles and Jernigan, 2010; The Neurobiology of Brain and Behavioral Development, 2018). Among others, neurulation gives rise to neural progenitors, neural crest, sensory placodes, and epidermis, all ectodermal derivatives (Haremaki et al., 2019). The appearance of these four lineages results from complex morphogenetic processes and several signaling activities, such as TGF-β inhibition and BMP4, Wnt, and FGF signaling pathways. The signaling molecules are represented already in non-linage committed iPSC from Huntington’s disease juvenile patients and mouse models, which show a range of molecular phenotypes such as MAPK, Wnt, and p53 pathways (Szlachcic et al., 2015, 2017).

Early human neurulation can be recapitulated in vitro by self-organizing neuruloids, containing cell populations present at the stage of neural tube closure in human development (days 21-25 post-fertilization) (Haremaki et al., 2019). Interestingly such Neuruloids generated from Huntington’s disease hESC demonstrated impaired neurogenesis resulting in aberrant rosette formation. In detail, HD 56Q neuruloids showed altered levels of Wnt/PCP pathway downregulation (for example, WNT5B and RSPO3 specific in neuroepithelium) and RHOB and RAB5C in the neural crest. In addition, decreased expression of cytoskeleton-associated genes and actin-myosin contraction (*EVL, MID1, RHOQ*, and *TMEM47*) could be observed and hint toward an impairment in the actin-mediated tissue organization mechanism during neurulation (Haremaki et al., 2019). In another recent study, one-third of gene changes in RNA-seq analysis on HD patient-derived iPSCs were involved in pathways regulating neuronal development and maturation. When these deregulated genes were mapped to stages of mouse striatal development, the profiles aligned mainly with earlier embryonic stages of neuronal differentiation (The HD iPSC Consortium, 2017). Moreover, sensory-motor network connectivity changes can be observed in the brains of HD patients, hinting at an effect of this PolyQ disease on brain connectivity (Pini et al., 2020).

During brain development, in a process called interkinetic nuclear migration coupled to cell cycle, neural progenitors keep the balance between the cell renewal of progenitors and their differentiation by controlling when and how many apical progenitor nuclei are exposed to proliferative versus neurogenic signals. Apical progenitors maintain their polarity through endocytosis and trafficking of glycans from the Golgi apparatus to the plasma membrane at the apical endfeet (Arai and Taverna, 2017). Interestingly, mislocalized expression of mHTT hinders both endosomal trafficking in apical progenitors, as well as the normal progression of cell cycle stages, leading to a shift towards more neural differentiation and away from proliferation (Barnat et al., 2020). Afterward, neuroepithelial cells start expressing glial genes and thereby begin a differentiation process into radial glial cells (RGCs). At this stage, cell migration starts to play a decisive role. Neuronal cells originating from the ventricular and subventricular zones start migrating outward in a radial fashion, using the RGCs as guideposts. Some subsets of RDGs eventually differentiate into intermediate, immature, and finally into mature neurons or astrocytes (Franco and Müller, 2013; The Neurobiology of Brain and Behavioral Development, 2018). Other cell populations migrate to the cortex during later developmental stages and include the microglia, which mostly use vessels for guidance into the forebrain. Recent reports point towards glia, particularly microglia, as essential players for cortical morphogenesis via regulation of brain wiring and interneuronal migration in the cortical wall (Silva et al., 2019).

Over time, successive layers of the cortical mantle form, and the progenitor cells are becoming more restricted in the cell types that they can construct. Furthermore, in this cellular maturation process, neural cells start to extend dendrites and an axon to form connections with other cells and become an integral part of a communication network (The Neurobiology of Brain and Behavioral Development, 2018).

In the prenatal stage of life, the further development of the brain also starts to depend on degenerative processes such as programmed cell death or apoptosis. These processes are initiated to remove the brain cells which have failed to make connections or have underutilized connections (Chan et al., 2002). Also, the underused synapses are eliminated in a process called synaptic pruning. In these stages of brain development, a transcriptional repressor complex of Ataxin1 and Capicua (ATXN1-CIC) regulates cell lineage specification and is involved in the regulation of cell proliferation (Ahmad et al., 2019). Loss of the ATXN1-CIC complex may have severe neurodevelopmental consequences, as conditional knockout of either Atxn1-Atxn1l or Cic in mice lead to a decrease of cortical thickness, hyperactivity and memory deficits (Lu et al., 2017). Indeed, loss or reduction of functional ATXN1 has been observed in patients with autism spectrum disorder and attention-deficit/hyperactivity disorder (Celestino-Soper et al., 2012; Di Benedetto et al., 2013), suggesting that loss of ATXN1-CIC complexes causes a spectrum of neurobehavioral phenotypes (Lu et al., 2017). Expanded CAG tracts in ATXN1 have been shown to stimulate the proliferation of postnatal cerebellar stem cells in SCA1 mice, which tend to differentiate into GABAergic inhibitory interneurons rather than astrocytes (Edamakanti et al., 2018). These hyperproliferating cells lead to a significantly increased number of GABAergic inhibitory interneuron synaptic connections, which in turn disrupt the proper cerebellar Purkinje cell function (Edamakanti et al., 2018). On the other hand, SCA2 patient fibroblast cells exhibit higher levels of caspase-8- and caspase-9-mediated apoptotic activation than those of healthy controls, which contributes to the pathophysiology of SCA2 (Wardman et al., 2020).

Also, the normal function of atrophin-1 and atrophin-2 proteins are related to the development and may be associated with regulation of cell polarity and transcriptional control of progenitors, which was reviewed previously (Shen and Peterson, 2009; Mannervik, 2014). Knockdown of Atn1 in neuronal progenitor cells (NPCs) in a rat led to severe aberrations in brain development. The study also highlighted ATN1 role as a direct target of the lysine-specific histone demethylase 1A (LSD1), which is known to have crucial developmental roles such as cortical neuronal migration or adult NPC proliferation (Zhang et al., 2014). Similarly, TATA Binding protein as the part of the TFIID complexes may control promoter elements can regulate of developmental transcription (Ohler and Wassarman, 2010). As a general transcription factor, TBP is, directly and indirectly, involved in numerous biological pathways. Studies confirmed many cellular processes impaired by mutant TBP via either gain of function or loss of function mechanisms, such as Notch signaling, TrkA signaling, Chaperone system, ER stress response, and muscle function (Yang et al., 2016).

Combined with the previously mentioned roles of HTT, ATXN1, ATXN2, ATXN3, ATN1, and TBP in transcription, translation, RNA metabolism, and ubiquitin-dependent protein quality control processes, a case can be made for the adverse effect of CAG tract extension on normal gene expression and protein regulation during neural development. Therefore, it can be proposed that other late-onset degenerative diseases may also be rooted in subtle developmental derailments. Deregulation of genes involved in cell migration, cell differentiation, maturation, synaptogenesis, and apoptosis can lead to severe neurodevelopmental disorders and may also contribute to the disease pathology of PolyQ diseases.

## 3. Different Brain Regions and Connections are Affected in Juvenile and Adult PolyQ diseases

PolyQ diseases affect a wide variety of brain regions, connections, and cell types in a heterogenic manner. In both juvenile- and adult-onset HD, the most affected cell types in the brain are striatal neurons (Tereshchenko et al., 2019). MRI data from JOHD cases show mostly cerebellar atrophy. The most substantial reduction in brain volume is observed in the caudate, putamen, as well as in globus pallidus and thalamus. Amygdala, hippocampus, and brainstem are slightly enlarged in HD patients (Hedjoudje et al., 2018). The significant difference between HD adults and children is seen in the cerebral cortex, which is mainly unaffected in children. Histopathological findings (Latimer et al., 2017) showed mild to moderate neuron loss in the brain tissue of adult-onset patients, while no significant loss of neocortical neurons was observed in JOHD. However, in JOHD patients, a significant neostriatal neuron loss and associated astrogliosis in the striatum were observed. In both disease onsets, HTT positive intranuclear and cytoplasmic neuronal inclusions can be found in the cerebral and striatum cortex.

SCAs present a broad range of dysfunctions in many brain structures such as the cerebellum, basal ganglia, brainstem, cerebral cortex, spinal cord, and peripheral nerves (Benton et al., 1998b). The most characteristic feature of general SCA1 pathology is the atrophy and loss of Purkinje cells from the cerebellar cortex. As SCA1 progresses, pathology is noted in other regions of the brain, including the deep cerebellar nuclei, especially the dentate nucleus, the inferior olive, the pons, and the red nuclei (Zoghbi and Orr, 2009). Juvenile onset is characterized by severe brainstem dysfunction in addition to the cerebellar symptoms. Subnormal cortical function and rapid progression of the disease are the most outstanding features (Zoghbi et al., 1988). The most affected cell types are Purkinje neurons in both early and late-onset cases (Zoghbi et al., 1988; Naphade et al., 2019). The MRI of children with very early-onset SCA2 (age from 7 to 17 months) revealed enlarged lateral ventricles, markedly small cerebellum and vermis, and associated atrophy involving the brainstem and both cerebral hemispheres. Moreover, increasing cerebral white matter loss, dysmyelination, pontocerebellar atrophy, and thinning of the corpus callosum was observed during SCA2 disease progression (Moretti et al., 2004; Ramocki et al., 2008; Paciorkowski et al., 2011; Vinther-Jensen et al., 2013; Singh et al., 2014). Histopathology findings in the cerebellar cortex showed a profound loss of Purkinje and granular neurons with severe attenuation of the molecular layer (Paciorkowski et al., 2011). Pathological examination of juvenile SCA3 patients has shown degeneration and mild gliosis of the substantia nigra, dentate, pontine and cranial nerve nuclei, anterior horns, and Clarke’s columns, with the consequent loss of fibers of the superior and middle cerebellar peduncles and spinocerebellar tracts (Coutinho et al., 1982). The most affected cell type in adult SCA3 are motor neurons (Naphade et al., 2019). However, in juvenile cases of SCA3, the dorsal root and trigeminal ganglia show severe nerve cell loss (Coutinho et al., 1982). A study by Wang et al. (Wang et al. 2010) showed that neurodegeneration in SCA6 also occurs in the spinal cord. Results of an autopsy of siblings with early-onset SCA6 revealed severe neurodegeneration in the cerebellum, dentate nucleus, and olivary nuclei (Wang et al. 2010). The most affected cell type in both adult and juvenile SCA6 are Purkinje cells (Wang et al., 2010; Naphade et al., 2019). Adult SCA7 is characterized by neural loss, mainly in the cerebellum and regions of the brainstem, particularly the inferior olivary complex (Holmberg, 1998). Juvenile cases present marked atrophy of both the cerebrum and cerebellum, ventricular dilation, as well as delayed myelination for age (Benton et al., 1998b). Other reports show diffuse volume reduction of the brain and increasing atrophy of the brainstem and cerebellum during SCA7 disease progression (Donis et al., 2015). The most affected cell types in SCA7 are retinal, cerebellar, and medulla oblongata neurons (Naphade et al., 2019). MRI data of 14 years old female with SCA17 showed prominent cerebellar atrophy accompanied by a dilatation of the fourth ventricle, and mild cerebral atrophy as well as dilatation of the lateral ventricles(Koide et al., 1999). It is familiar with neuroimaging studies of a family with age at onset range from very early to adult-onset that showed cerebral and cerebellar atrophy in all patients (Maltecca et al., 2003). The most affected cell types in SCA17 are Purkinje, medium spiny cortical, and dopaminergic neurons (Naphade et al., 2019).

DRPLA is characterized by severe neuronal loss in the dentatorubral and pallidal-subthalamic nucleus (corpus Luysii). In the juvenile type, presenting with PME syndromes, degeneration of the globus pallidus was observed to be more severe than that of the dentate nucleus. Atrophy of the brainstem and spinal cord was noticed as mild (Takeda and Takahashi 1996). MRI data of children with DRPLA also showed severe atrophy of the cerebrum and cerebellum, delayed myelination, and thin corpus callosum (Shimojo et al. 2001). In general, juvenile-onset can be characterized by more marked pallidoluysian degeneration than dentatorubral degeneration, which is opposite to late-adult onset degeneration pattern (Yamada, 2010). Histochemistry revealed nonspecific cerebral atrophy and mild neuronal loss with gliosis in the cerebral cortex (Hayashi et al. 1998; Tsuchiya et al. 1998). The most affected cell types in DRPLA are striatal medium spiny neurons and pallidal neurons (Naphade et al., 2019).

## 4. Review of Juvenile- and Adult-Onset HD and other PolyQ diseases: Deregulated Genes Overlap and GO Terms Over-Representation Analysis

To obtain a broader view of the role of the very long CAG repeats and very long polyQ tracts in proteins in early brain development, we collected published transcriptomic data from human juvenile- and adult-onset HD (An et al., 2012; Feyeux et al., 2012; HD iPSC Consortium, 2012a; Chiu et al., 2015a; Ring et al., 2015; Nekrasov et al., 2016a; The HD iPSC Consortium, 2017; Mehta et al., 2018a; Świtońska et al., 2019a; Al-Dalahmah et al., 2020; Smith-Geater et al., 2020a) and also published RNA-seq or microarray data from different PolyQ mouse models (Suzuki et al., 2012; Aikawa et al., 2015; Agostoni et al., 2016; Pflieger et al., 2017; Driessen et al., 2018; Hervás-Corpión et al., 2018; Malik et al., 2019; Liu et al., 2020; Stoyas et al., 2020). The published mice data from SCA1, SCA2, SCA6, SCA7, SCA17, and DRPLA were originally collected from different brain regions, however, data from SBMA mice were collected from primary motor neurons in the spinal cord. We first focused on the publications with human data where the main aim was to compare genes dysregulated in two types of HD onset in a more detailed way. The analysis of overlapping deregulated genes (DEGs) between diseases was created and visualized with R software 3.6.3 (R Core Team 2018) and its three packages: UpsetR (Conway et al., 2017), ComplexHeatmap (Gu et al., 2016), and VennDiagram (Chen and Boutros, 2011). GO terms over-representation analysis was conducted in Cytoscape (Shannon et al. 2003) and its ClueGO app (Bindea et al., 2009, 2013). An overview of data from all papers included in the analysis can be found in Supplementary Table S1. Transcriptomic data were retrieved from the Gene Expression Omnibus (GEO) repository, if possible, or from the supplementary material provided with the original publication. A cut-off of *p value* < 0.05 was considered as significant. In two papers, with much higher number of identified genes, we set a cut-off of p value < 0.001 (HD iPSC Consortium, 2012a; The HD iPSC Consortium, 2017).

### 4. Previously Published Transcriptomic Data Show Molecular Downregulation in Juvenile-Onset Human HD and Highlights Organism Morphogenesis, Neurodevelopment, and Synaptic Transmission

First, we assessed the overlap of DEGs between different cell types and between different types of HD disease onset (Supplementary Figure1 and 2). We focused on data from embryonic stem cells (ESC), induced pluripotent stem cells (iPSC), neural stem cells (NSC), and neurons. With such a collection, we were able to check whether there are genes downregulated in HD from the very beginning and at the same time through the whole “neurodifferentiation axis”. The analyses revealed two genes shared between iPSC, NSC, and neurons in data from JOHD, *TBX15*, and *HOXB6* (Supplementary Figure 1A-B and Supplementary Table S1). These two genes encode transcription factors that regulate a variety of developmental processes. We identified 12, and 22 genes shared between iPSC and NSC with neurons, respectively, in JOHD (Supplementary Figure 1A-B and Supplementary Table S1). The firstly mentioned 12 genes are again connected in the majority with the regulation of transcription. The NSC/neurons shared genes are involved in developmental biology and particularly on embryonic skeletal system morphogenesis. When it comes to adult-onset HD, apart from a small group of 11 genes shared between ESC and NSC, we didn’t identify genes downregulated in every cell type (Supplementary Figure 1C and Supplementary Table S1). Altogether, the created Venn diagrams highlight the fact that in JOHD, molecular processes and genes downregulated on very early stages of organism development may have a direct impact on later brain and neurons formation, hence resulting in a much earlier disease onset. The UpSetR diagram did not show much of an overlap of downregulated genes between juvenile and adult HD (Supplementary Figure 1A). Nonetheless, 27 genes were identified to be DEGs in neurons obtained from both disease onset type (Supplementary Figure 1D-E). Those are involved, among others, in the cerebral cortex GABAergic interneuron differentiation, which aberration leads to an imbalance between excitatory and inhibitory signaling, affecting motor and cognitive processes during HD pathogenesis (Hsu et al., 2018). We also analyzed which biological processes include genes downregulated only in juvenile or only in adult HD. This resulted in a big cluster of various early neurodevelopmental processes, organism morphogenesis, and signal transduction for JOHD (Figure 1), which was not the case for adult HD. Besides some neuronal GO terms connected with genes downregulated in adult HD, no obvious cluster of connected processes was identified. Particularly interesting were the four papers with transcriptomic data on human juvenile-onset HD neurons and four articles concerning human adult-onset HD neurons, which we compared (HD iPSC Consortium, 2012b, 2012b; Chiu et al., 2015b; Nekrasov et al., 2016b; Mehta et al., 2018b; Świtońska et al., 2019b; Al-Dalahmah et al., 2020; Smith-Geater et al., 2020b). A total of 27 downregulated and 48 upregulated genes in neurons were found to be shared between juvenile-onset and adult-onset HD (Supplementary Figure 1D & 2D). A total of 758 downregulated and 632 upregulated genes in neurons were found to be unique for juvenile-onset HD, and an additional 108 downregulated and 451 upregulated genes in neurons were unique to adult-onset HD (Supplementary Figure 1D & 2D). A full list of common and uniquely deregulated genes can be found in the supplemental data of this work (Supplementary Table S1).

**FIG 1.**
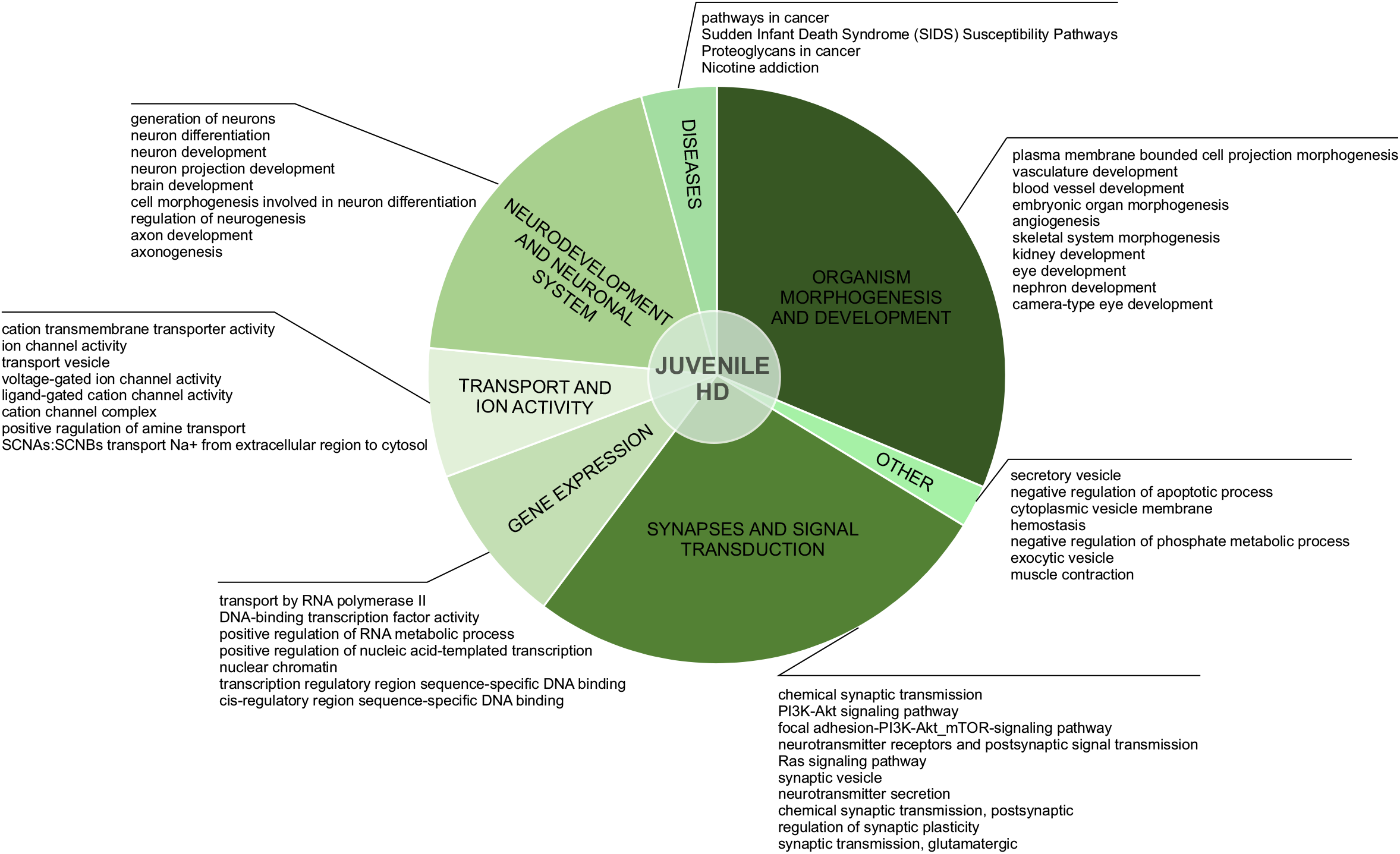
Uniquely downregulated DEGs in JOHD are involved in developmental processes, organism morphogenesis, and signal transduction. A pie-chart with ClueGO analysis of genes downregulated only in neurons from juvenile-onset HD patients was used to visualize the biological processes they are involved in. Top dysregulated processes for each bigger cluster were listed.

After the assessment of gene overlap, we performed pathway analysis with ClueGO app (Cytoscape). We found that the DEGs uniquely downregulated in juvenile-onset HD neurons are significantly involved in developmental processes, such as Dopaminergic Neurogenesis (PW:0000394), Differentiation Pathway (WP2848), spinal cord development (GO:0021510), Neuronal System (R-HSA-112316.7), Neural Crest Differentiation (WP2064), presynaptic active zone assembly (GO:1904071), anterior/posterior axon guidance (GO:0033564, metencephalon development (GO:0022037), Potassium Channels (WP2669), and DNA-binding transcription activator activity, RNA polymerase II-specific (GO:0001228) Besides developmental processes, a substantial subset of the uniquely downregulated genes in JOHD-derived neurons is involved in synaptic processes, regulation of synaptic transmission, glutamatergic (GO:0051967 and GO:0051968), Cholinergic synapse (GO:0098981), neurotransmitter secretion (GO:0007269), axon terminus (GO:0043679), positive regulation of dopamine secretion (GO:0033603), regulation of neuronal synaptic plasticity (GO:0048168), and regulation of dendrite morphogenesis (GO:0048814). In Supplementary Table S2, we present a list of the most significantly involved pathways in uniquely downregulated DEGs in JOHD or adult-onset HD, grouped by biological processes, and highlight the input genes found in those pathways. The GO terms unique to neurons of adult-onset HD patients suggest a more developed, more mature cellular expression pattern compared to the juvenile-onset HD.

Inspired by transcriptomic data generated by Haremaki and colleagues (Haremaki et al. 2019) we decided to extend our bioinformatic study with one additional comparative analysis. As previously mentioned, Haremaki and colleagues succeeded in recapitulating human neurulation by generating neuruloids harboring neural progenitors, neural crest, sensory placode and epidermis. These self-organizing structures provide a great opportunity to study the developmental aspects of many human diseases, especially HD. Having the insight into single-cell transcriptomics from healthy and HD neuruloids, we decided to compare these data with the ones collected for our comparative study. We compared down- and upregulated genes from our cohort to each group of markers specific to a particular cell population identified in scRNA-seq of healthy neuruloids, neuroepithelial identity NE1 and NE2, neurons, skin, neural crest (NC), placode and U1 neurons, and also to a list of differentially expressed genes in NE and NC populations in HD neuruloids (Supplementary Table S4). We identified a significant number of genes shared between markers for neuruloid neurons population and downregulated genes in stem cell-derived neurons in juvenile-onset HD (Supplementary Table S4). This is coherent with GO term over-representation analysis and again highlights the great downregulation of crucial genes and thus many biological processes during the very early neurogenesis.

### 4. HD and SCA1 Seems to Have More Common Transcriptionally Dysregulated Genes Than Other PolyQ diseases in Mice

Being rare diseases, more abundant data can be drawn from mice models of PolyQ diseases. An extensive review of polyQ mouse models can be found in the works of Figiel et al. and Switonski et al. (Figiel et al., 2012; Switonski et al., 2012). The high CAG repeat numbers is needed in polyQ mouse models to express a disease phenotype, therefore they may be considered as polyQ models of juvenile-onset type. Therefore, the second data collection for this bioinformatic review was derived from nine publications concerning mouse brain transcriptomics in several PolyQ diseases, such as HD, SCA1, SCA2, SCA6, SCA7, SCA17, DRPLA, and SBMA (Suzuki et al., 2012; Aikawa et al., 2015; Agostoni et al., 2016; Pflieger et al., 2017; Driessen et al., 2018; Hervás-Corpión et al., 2018; Malik et al., 2019; Liu et al., 2020; Stoyas et al., 2020). (Supplementary Table S1). After adjusting *p-value* cut-off, the following number of genes was collected: 697 downregulated and 167 upregulated DEGs in HD and respectively 643 and 144 in SCA1, 134 and 80 in SCA2, 493 and 349 in SCA6, 64 and 27 in SCA7, 246 and 187 in SCA17, 250 and 162 in SBMA, 225 and 318 in DRPLA (Figures 2 and 3 and Supplementary Table S1). The largest subset of commonly shared DEGs were 87 downregulated genes common between HD and SCA1 (Figure 2B, Supplementary Table S3). ClueGo analysis revealed the involvement of DEGs in Amphetamine addiction (KEGG hsa05031), Opioid signaling (WP1978), neuronal cell body membrane (GO:0032809), and integrin cell surface markers (WP1833) (Figure 2C, Supplementary Table S5). SBMA stood out as the least common of the PolyQ diseases, with 235 out of 250 downregulated and 152 out of 162 upregulated genes being uniquely expressed in SBMA only (Figure 2A and 3A).

**FIG 2.**
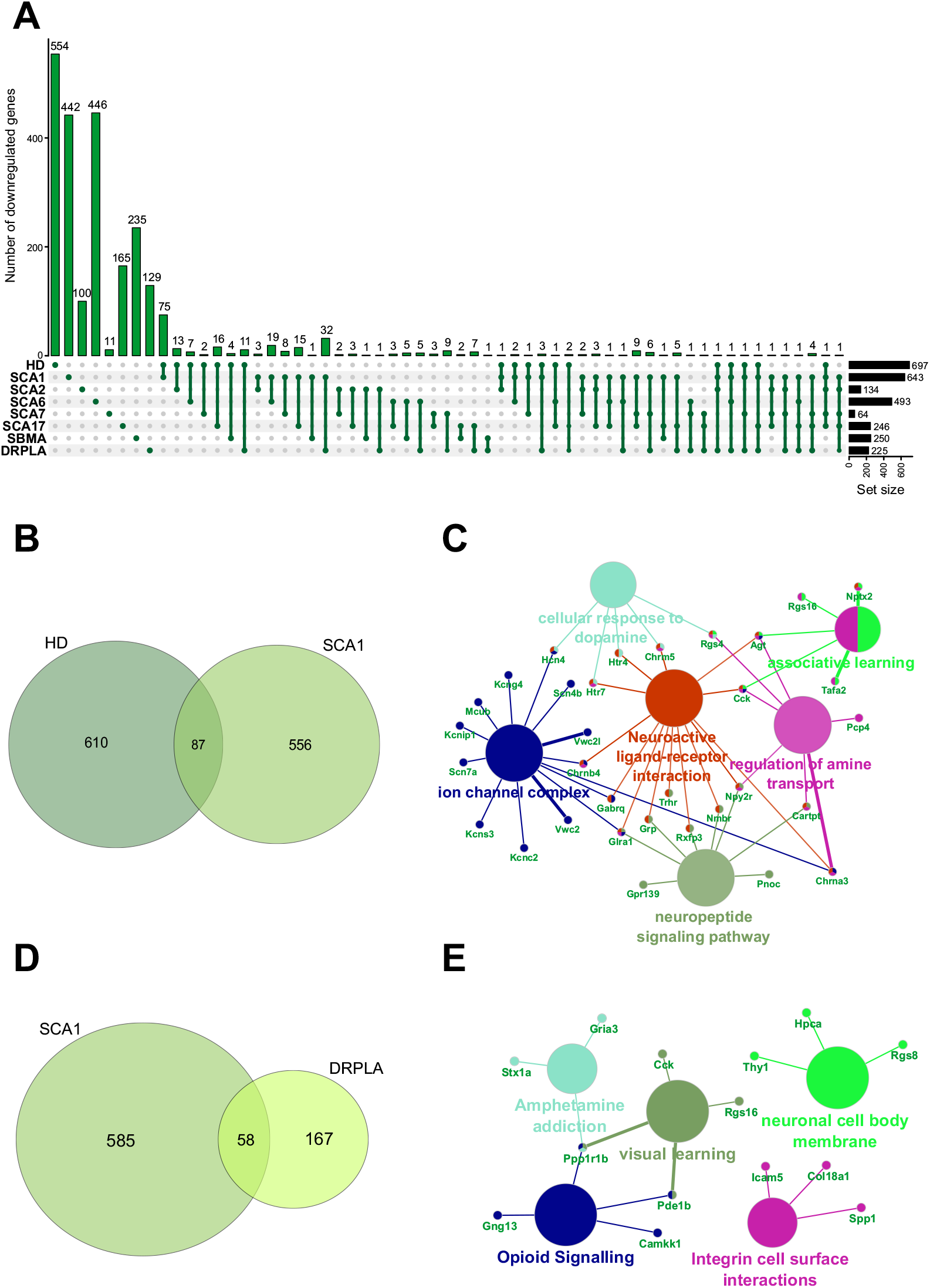
Overlap of significantly downregulated genes from mice transcriptomic data from different PolyQ diseases. (A) UpsetR analysis was used to see the overlap between downregulated genes identified in different PolyQ diseases. Venn diagrams visualizing the overlap between downregulated genes in HD and SCA1; (B) and for overlapping genes downregulated in SCA1 and DRPLA (D). ClueGO analysis was used to visualize the biological processes in which the commonly downregulated genes between HD and SCA1 (C) and between SCA1 and DRPLA (E) are involved.

**FIG 3.**
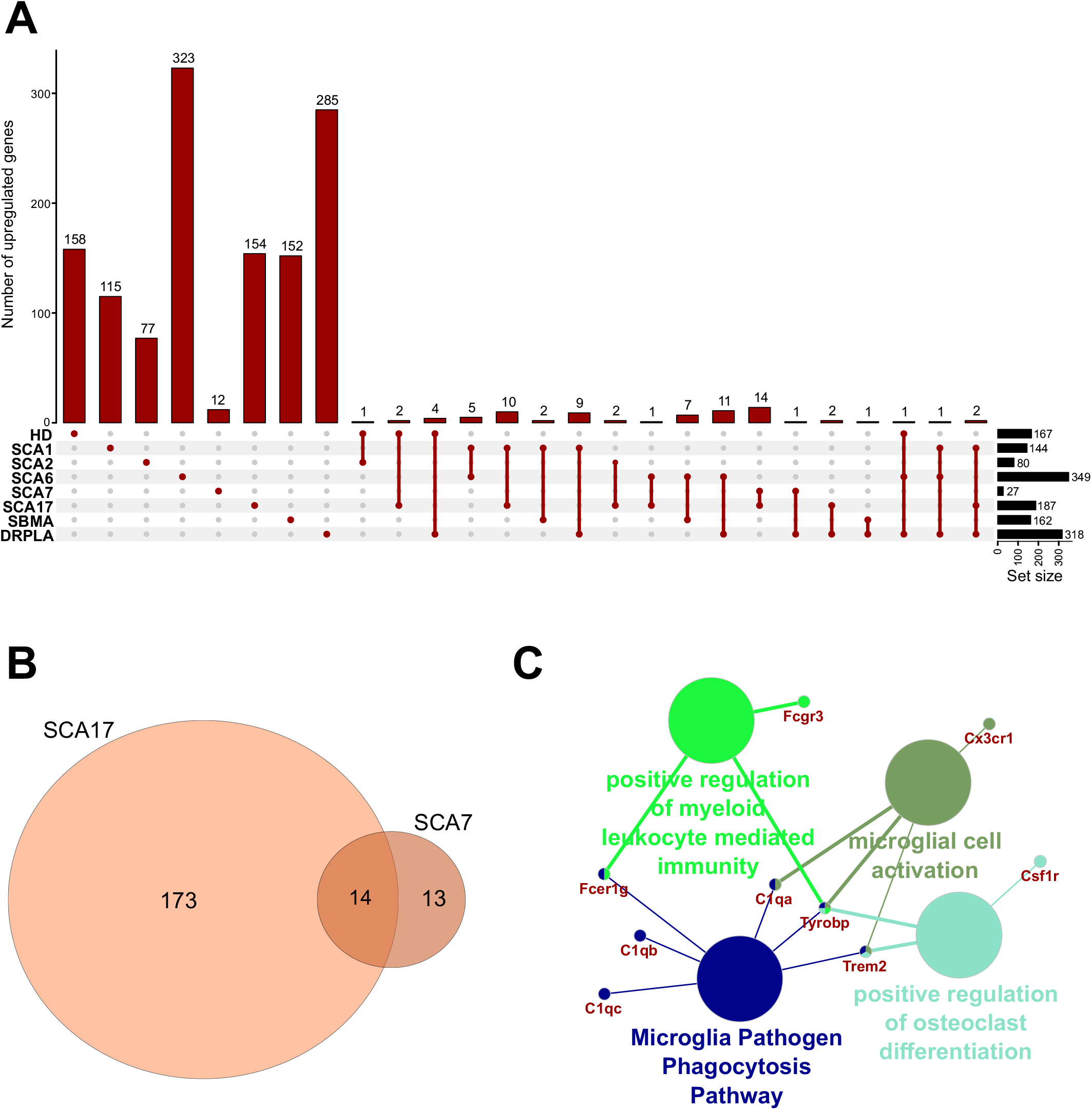
Overlap of significantly upregulated genes from mice transcriptomic data from different PolyQ diseases. (A) UpsetR analysis was used to see the overlap between upregulated genes identified in different PolyQ diseases. (B) Venn diagrams visualizing the overlap between genes upregulated in SCA7 and SCA17. (C) ClueGO analysis for genes commonly upregulated in SCA7 and SCA17.

Two genes were shared between five of the PolyQ diseases (G Protein Subunit Gamma 13 (*Gng13)* in SCA1, 2, 7, 17 and DRPLA, and Glutamate receptor delta two interacting protein (*Grid2ip*) in HD, Sca1, 2, 7, and 17). *Gng13* encodes the gamma subunit of heterotrimeric G proteins, which are signal transducers for the 7-transmembrane-helix G protein-coupled receptors (Li et al., 2006). Grid2ip is a Purkinje cell-specific postsynaptic protein, where it may serve to link Glutamate receptor delta 2 (GRID2) with the actin cytoskeleton and various signaling molecules. GRID2 has been reported to play crucial roles in synaptogenesis and synaptic plasticity and may control GRID2 signaling in Purkinje cells (Matsuda et al., 2006). Other notable DEGs are Regulator Of G Protein Signaling 8 (*Rgs8*), Regulator Of G Protein Signaling 16 (*Rgs16*), and Purkinje Cell Protein 4 (*Pcp4*) commonly deregulated in HD, SCA1, DRPLA, and either SCA6 (*Rgs16*) or SCA7 (*Rgs8* and *Pcp4*). These DEGs are all involved in calmodulin-binding, which acts as part of a calcium signal transduction pathway and has roles in cellular mechanisms including metabolism, synaptic plasticity, nerve growth, smooth muscle contraction (Hyman and Pfenninger, 1985; Xia and Storm, 2005; Kleerekoper and Putkey, 2009; Mouton-Liger et al., 2011; Wang and Putkey, 2016)

Finally, several Cerebellin (*Cbln1*, 2, 3 and 4), Matrix Metalloproteinases (*Mmp8*, 9, 16, 17, and 20), and Collagen (*Col5a1, Col6a4, Col11a1, Col18a1, Col20a1, Col25a1*)) isoforms are downregulated in compared PolyQ diseases. While no commonly deregulated isoform was found, the downregulation of these proteins is important for synaptic activity and the modulation of the extracellular matrix, further hinting to an important role of WM alterations in PolyQ diseases.

## 5. Discussion and Concluding Remarks

Although the juvenile and infantile forms make up a minority of PolyQ disease cases, the early onset makes these diseases an example of neurodevelopmental disorders. Indeed, the results of our bioinformatic study of the available transcriptomic data reveal that uniquely dysregulated genes in juvenile-onset HD neurons are involved in several (neuro)developmental pathways leading to early symptoms in patients. Our group and others have previously demonstrated a neurodevelopmental component in HD pathogenesis, and further exciting evidence was delivered only very recently (Kubera et al., 2019; Barnat et al., 2020). Moreover, HTT has an impact on the cortical volume and brain connections, leading to higher general intelligence (IQ) in people with larger (sub-disease) PolyQ repeats (Lee et al., 2017, 2018a). An increasing number of studies created a body of evidence for transcriptional modulators of polyglutamine tracts not only in HD but also in other PolyQ diseases, like SCAs, mentioned in this manuscript (Paulson et al., 2017; Buijsen et al., 2019).

Our analysis combines numerous data sets on polyQ transcriptomics into one collection and demonstrates several neurodevelopmental transcriptomic commonalities to the diseases. There are genes unique in JOHD neurons and individual genes that are downregulated in 4 or more of the independent PolyQ diseases mouse models. The genes were involved in neural growth, synaptogenesis, and synaptic plasticity, and extracellular matrix remodeling, suggesting a critical role of brain connections and WM changes roles in PolyQ disease pathology. *HTT, ATN1, TBP*, and Ataxins have previously been identified as transcriptional regulators (Benn et al., 2008; Kumar et al., 2014; Gao et al., 2019) therefore, our results are in agreement with the previously formulated hypothesis that transcriptional dysregulation is a solid feature of several polyglutamine diseases (Helmlinger et al., 2006).

Polyglutamine diseases are relatively rare, and therefore, only a limited number of publications with transcriptomic data were available for our comparative study. Therefore more transcriptomic research in PolyQ disease is needed to understand better the mechanistic aspects of the disease pathology. Moreover, studies that will focus on the unique differences between juvenile- and adult-onset would be of interest, as the longer CAG repeat mutations augment the transcriptional potential of the affected protein, which may leading to compromised of neurodevelopment.

## AUTHOR CONTRIBUTIONS

K.Ś-K., B.K., J.D and M. Figiel wrote the manuscript. K.Ś-K performed all bioinformatics associated with R software and ClueGO analyses of data. All authors read and approved the final manuscript. M. Figiel was responsible for concept of this review and for obtaining funding.

## ACKNOWLEDGMENTS

This work was supported by the grant from the National Science Centre (grant number 2018/31/B/NZ3/03621).

## ADDITIONAL INFORMATION

The authors declare no competing or financial interests.

## SUPPLEMENTARY DATA

**SUPPLEMENTARY FIGURE S1**. Analysis of Juvenile- and adult Huntington disease transcriptomic data demonstrates mostly specific sets of downregulated genes for each type of onset. (A) UpsetR graph showing the intersection between genes identified in the different HD cell types. Venn diagrams were used to visualize the overlap between genes from juvenile HD iPSC, NSC, and neurons (B), for genes from adult HD iPSC, NSC, and neurons (C), and for genes from juvenile and adult neurons (D). Interestingly, although both the juvenile and adult neurons contain a mutation in HTT, their transcriptomic dysregulated genes vastly differ, showing just 27 genes in common. These commonly downregulated genes are visualized with a CluGO plot (E).

**SUPPLEMENTARY FIGURE S2**. Analysis of Juvenile- and adult Huntington disease transcriptomic data demonstrates mostly specific sets of upregulated genes for each type of onset. (A) UpsetR graph showing the overlap between genes identified in the different HD cell types. Venn diagrams were used to visualize the overlap between genes from juvenile HD iPSC, NSC, and neurons (B), for genes from adult HD iPSC, NSC, and neurons (C), and for genes from juvenile and adult neurons (D). Similar to Figure S1D, the juvenile HD and adult HD neurons vastly differ in dysregulated genes, showing only 48 genes in common. These commonly downregulated genes are visualized with a CluGO plot (E).

**SUPPLEMENTARY TABLE S1**. Transcriptomic data included in the comparative bioinformatic study. Data were retrieved from the Gene Expression Omnibs (GEO) repository, if possible, or from the supplementary material provided with the publication. A cut-off of p-value < 0.05 was considered as significant. In two publications, with a greater number of identified genes, we set a cut-off of p-value < 0.001.

**SUPPLEMENTARY TABLE S2**. Biological processes, molecular function, and cellular components ClueGO analysis for genes downregulated in neurons from juvenile-onset HD patients (stem cell-derived or collected post-mortem). Top downregulated processes were visualized in Figure 3.

**SUPPLEMENTARY TABLE S3**. Biological processes, molecular function, and cellular components ClueGO analysis for common transcriptionally downregulated genes in HD and SCA1 mice.

**SUPPLEMENTARY TABLE S4**. Comparison analysis between scRNA-seq data from neuruloid paper (Haremaki et al., 2019) and human data collected for our comparative study. We compared down- and upregulated genes from our cohort to each group of markers specific to a particular cell population identified in scRNA-seq of healthy neuruloids, neuroepithelial identity NE1 and NE2, neurons, skin, neural crest (NC), placode and U1 neurons, and also to a list of differentially expressed genes in NE and NC populations in HD neuruloids.

