## Supplementary Figure S1. for "Juvenile Huntington’s Disease and Other PolyQ diseases, Update on Neurodevelopmental Character and Comparative Bioinformatic Review of Transcriptomic Data"

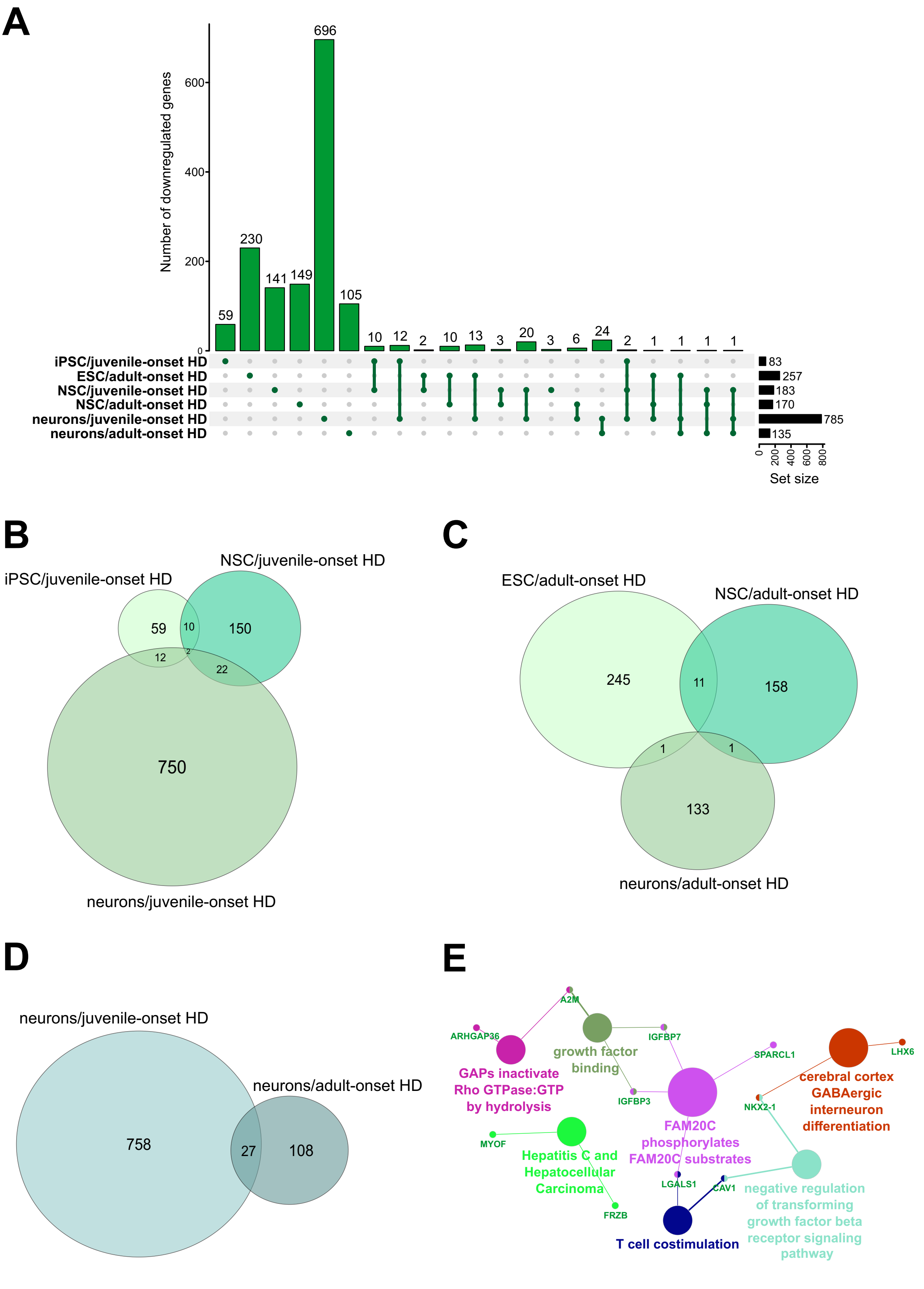


**SUPPLEMENTARY FIGURE S1.** Analysis of Juvenile- and adult Huntington disease transcriptomic data demonstrates mostly specific sets of downregulated genes for each type of onset. (A) UpsetR graph showing the intersection between genes identified in the different HD cell types. Venn diagrams were used to visualize the overlap between genes from juvenile HD iPSC, NSC, and neurons (B), for genes from adult HD iPSC, NSC, and neurons (C), and for genes from juvenile and adult neurons (D). Interestingly, although both the juvenile and adult neurons contain a mutation in HTT, their transcriptomic dysregulated genes vastly differ, showing just 27 genes in common. These commonly downregulated genes are visualized with a CluGO plot (E).
