## Supplementary Figure S2. for "Juvenile Huntington’s Disease and Other PolyQ diseases, Update on Neurodevelopmental Character and Comparative Bioinformatic Review of Transcriptomic Data"

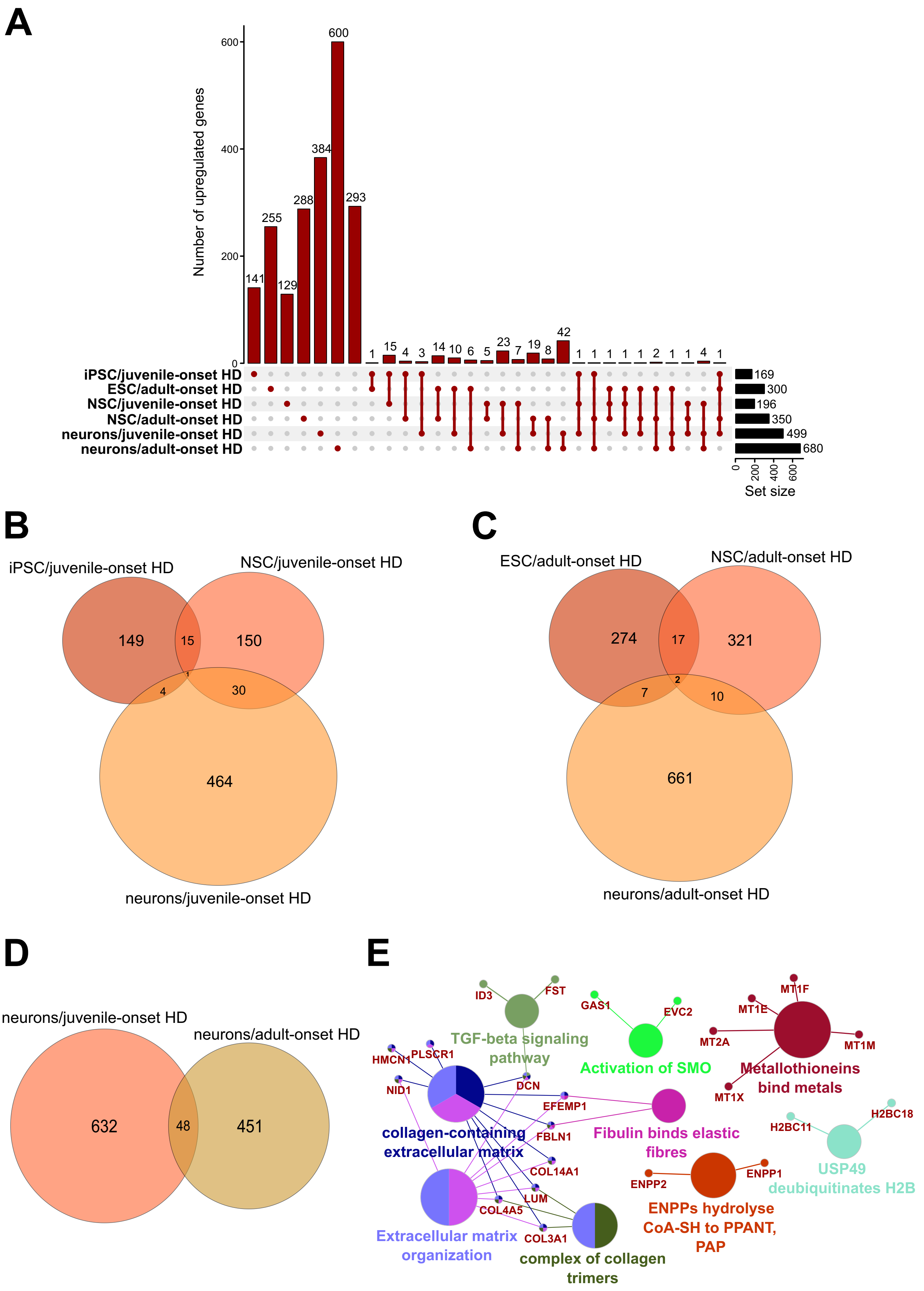


**SUPPLEMENTARY FIGURE S2.** Analysis of Juvenile- and adult Huntington disease transcriptomic data demonstrates mostly specific sets of upregulated genes for each type of onset. (A) UpsetR graph showing the overlap between genes identified in the different HD cell types. Venn diagrams were used to visualize the overlap between genes from juvenile HD iPSC, NSC, and neurons (B), for genes from adult HD iPSC, NSC, and neurons (C), and for genes from juvenile and adult neurons (D). Similar to Figure S1D, the juvenile HD and adult HD neurons vastly differ in dysregulated genes, showing only 48 genes in common. These commonly downregulated genes are visualized with a CluGO plot (E).
